# Can fallopian tube anatomy predict pregnancy and pregnancy outcomes after tubal reversal surgery?

**DOI:** 10.1101/616508

**Authors:** Rafael S. de Souza, Gary S. Berger

## Abstract

We conducted this study to determine if postoperative fallopian tube anatomy can predict the likelihood of pregnancy and pregnancy outcomes after tubal sterilisation reversal.

We built a flexible, non-parametric, multivariate model via generalised additive models to assess the effects of these tubal parameters observed during tubal reparative surgery: tubal lengths; differences in tubal segment location and diameters at the anastomosis sites; and, fibrosis of the tubal muscularis.

Age and tubal length (in that order) are the primary features determining the likelihood of pregnancy. For pregnancy outcomes, age is the primary predictor of miscarriage, but tubal length is the most influential predictor of the odds of birth and ectopic pregnancy. Segment location and diameters contribute slightly to the odds of miscarriage and ectopic pregnancy, whereas fibrosis has little apparent effect.

This study is the first to show that a statistical learning predictive model based on fallopian tube anatomy can predict both pregnancy and pregnancy outcome probabilities after tubal reversal surgery.

## Introduction

Female sterilisation is the most common contraceptive method worldwide^1^. Approximately 9.5 million US women rely on female sterilisation for birth control^2^. While sterilisation is intended to be permanent, postmodern society has experienced paradigmatic behavioural changes with increased rates of divorce and remarriage^3^. In this context, many sterilised women have expressed regret and wish to have their fertility restored. The frequency rate of the request can be as high as 14.3%^4^.

Despite reproductive endocrinologists’ preference to perform *in vitro* fertilisation, fertility restoration by tubal reparative surgery is a better choice for some women. Preoperative counselling based on age and method of sterilisation can estimate the likelihood of pregnancy and pregnancy outcomes after tubal anastomosis^5^. After reparative surgery, patients usually ask for information about their fallopian tubes and how it affects their prognosis. This study aims to answer this question.

This report presents a model for predicting pregnancy and pregnancy outcome probabilities based on a woman’s age and tubal anatomy observed during tubal reversal surgery.

## 1 Data acquisition and pre-processing

This section introduces the dataset utilised in this study, the pre-processing of our data, and the statistical models employed for the analysis of pregnancy and pregnancy outcomes.

### 1.1 Study design and population

This study makes use of a dataset from an observational study of women who had tubal reconstructive surgeries. The operations were performed in an outpatient surgical centre in Chapel Hill, NC, USA, from January 2000 to June 2013. The surgical and study methods have been described in detail previously^5^. The University of North Carolina at Chapel Hill IRB and Office of Human Research Ethics gave this study exempt status (IRB Number 14-1783) as a quality improvement study, meaning that written consent was not required.

Tubal anatomy was assessed at surgery for both segments joined at the anastomosis site for right and left fallopian tubes individually. The measurements included: length; location (interstitial, isthmic, ampullary, infundibular, or fimbrial); diameter; and presence of fibrosis. Location differences were recorded numerically. If the segments were identical (such as isthmic-isthmic or ampullary-ampullary) the location difference was 0. Diameters of the anastomosed segments were categorised as similar, somewhat dissimilar, or dissimilar. Fibrosis of the tubal muscularis, assessed visually and by palpation, was recorded as none, mild, moderate, or severe.

Among all women in the tubal surgery database, we retrieved information from the those who had bilateral tubotubal anastomosis with complete information about all fallopian tube anatomic parameters and at least one year of follow-up after surgery. These 5682 women comprise the study population for the present analysis.

The women’s ages at the time of reconstructive surgery ranged from 20 to 51 years. The distribution of age groups and sterilisation methods within each age group are shown in Fig. 1. In this study population, 19.3% were younger than 30 yrs, 40.8% were 30-35 yrs, 30% were 35-40 yrs, and 9.7% were 40 yrs of age or older. Across all age groups, the most common method of sterilisation was ligation/resection, followed in order by coagulation and by mechanical methods (rings or clips).

**Figure 1.**
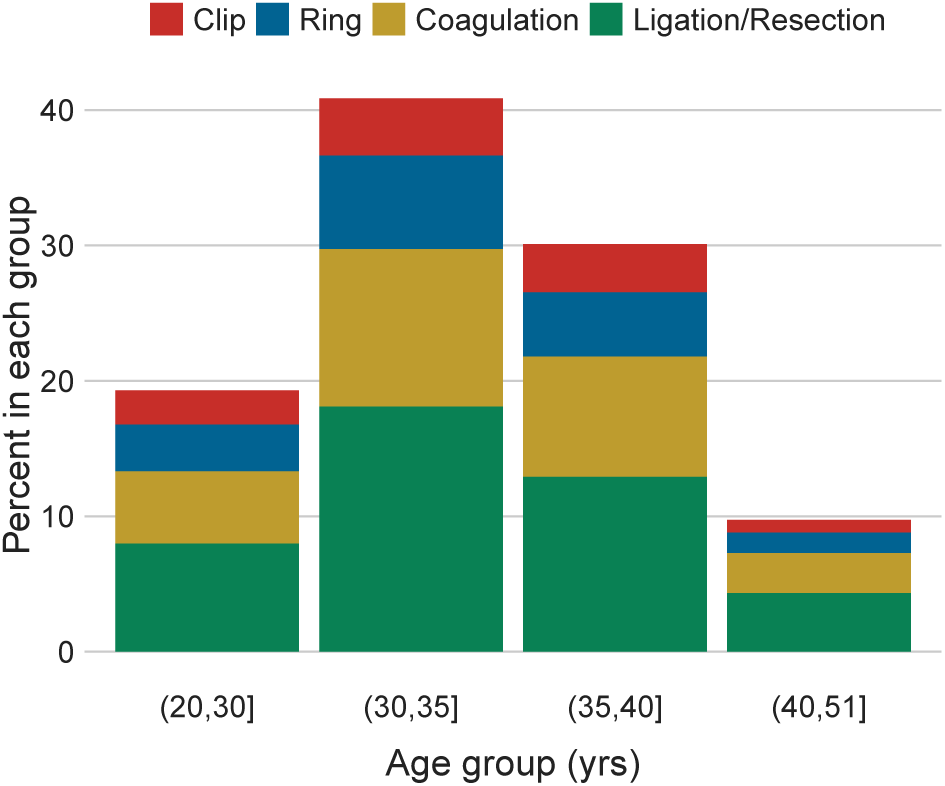
Age group distribution of 5682 women in which 19.3% were younger than 30 yrs, 40.8% between 30-35 yrs, 30% between 35-40 yrs, and 9.7% older than 40 yrs. Sterilisation methods are colour-coded, with ligation/resection representing the most common method across all ages.

### 1.2 Feature selection

For every record, we extracted the woman’s age in combination with the following anatomic properties for both left and right fallopian tubes: tubal length after anastomosis, specific tubal segments rejoined, diameters of the two segments at the anastomosis site, and fibrosis of the tubal muscularis.

Which one of a woman’s fallopian tubes results in a given pregnancy is unknown. Therefore, we emulated the randomness of the choice between left and right sides by attributing to each woman either left or right tubal anatomic properties following a Bernoulli process with 50% probability to chose one side or another. A Bernoulli distribution describes a sequence of independent experiments (trials) each of which has only two possible outcomes {0, 1}. In the specific case of interest here, one can think of the choice between left and right sides as binary data which is either left = 0, or right = 1. Fig. 2 displays the distribution of tubal anatomic properties for the study population in term of left and right features. Simple visual inspection does not reveal any directional bias towards a particular side.

**Figure 2.**
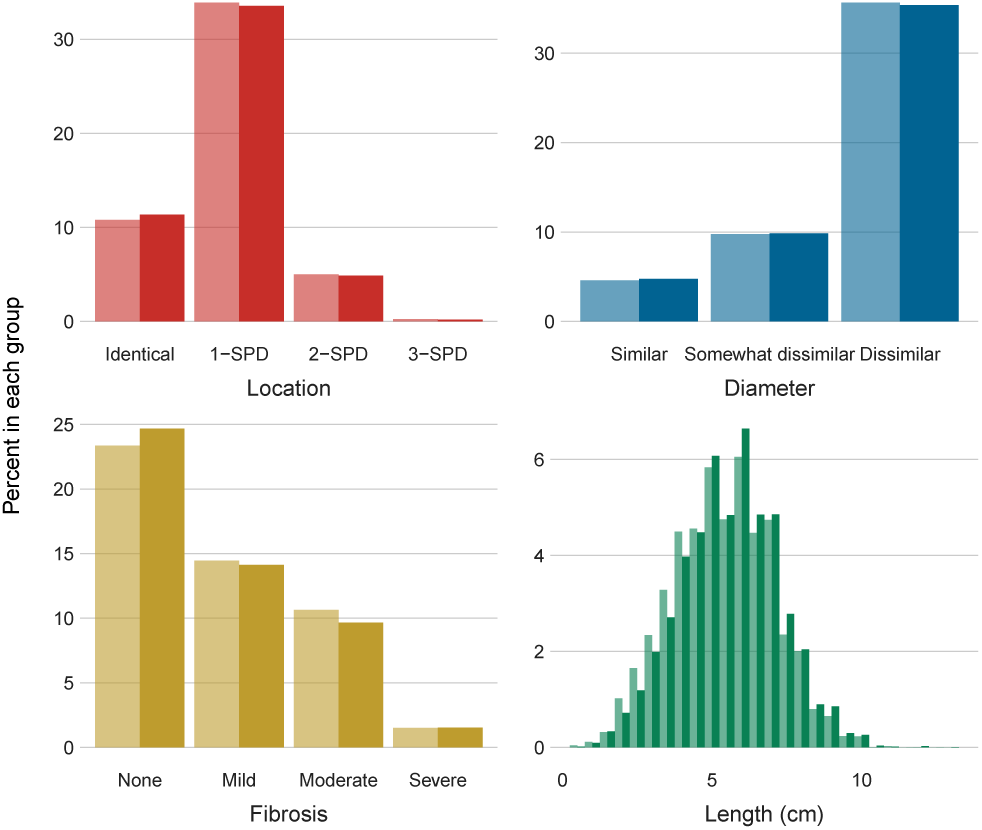
Percent in each group for anatomic properties: Location differences, upper left panel; Diameter differences, upper right panel; Fibrosis, lower left panel; Tubal length, lower right panel. In each panel the lighter bars represent the left tube, and the darker bars the right tube. For Location, SPD stands for segment position difference.

Every record was then represented as a 5-dimensional feature vector with 2 numeric properties (age, tubal length) and 3 categorical properties (location differences, diameter differences, extent of fibrosis). For the response variable, there are two parts of the analysis. The first concerns pregnancy prevalence and retrieves binary information: the woman became pregnant or not. The second part exploits each pregnancy outcome: birth, miscarriage, ectopic, or ongoing at time of last contact.

Fig. 3 shows the distribution of anatomic properties in terms of pregnancy occurrence. The median tubal length is slightly larger for women who became pregnant (5.5 cm) than those who did not (5 cm). Likewise, there appears to be a preference towards anastomosis of segments of identical location or 1-segment position difference and for none or mild fibrosis of the tubal muscularis among the pregnant women. Diameter differences do not show any visually detectable bias between pregnant and non-pregnant women.

**Figure 3.**
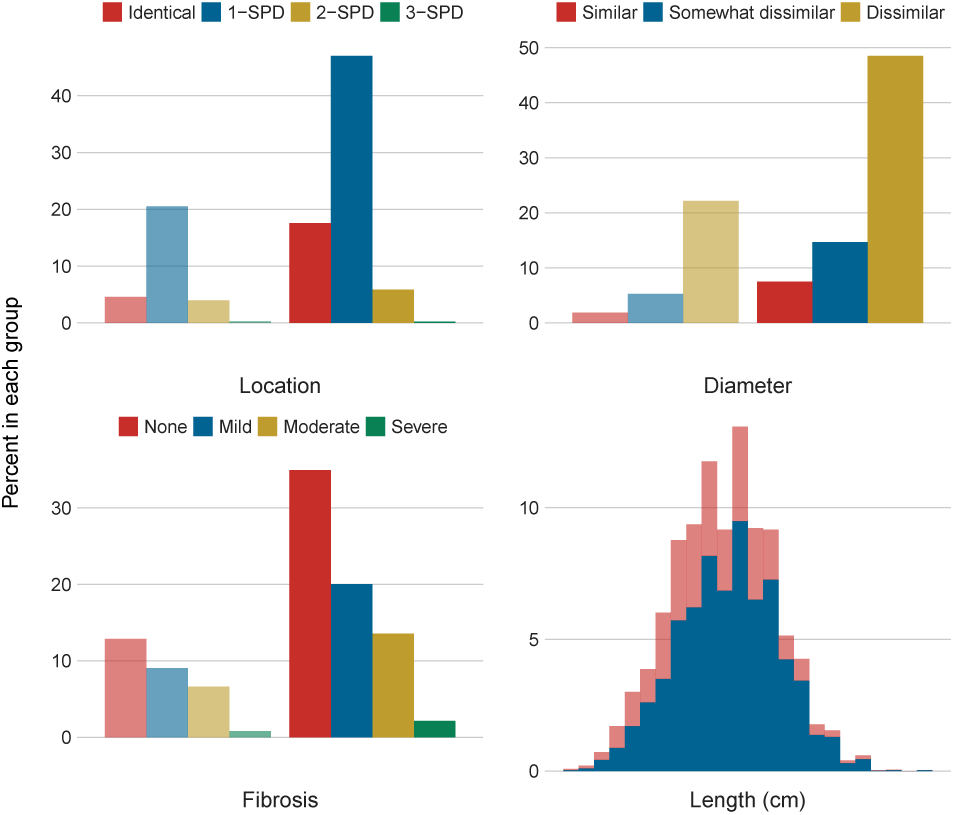
Percent in each group for anatomic properties: Location differences, upper left panel; Diameter differences, upper right panel; Fibrosis, lower left panel; Tubal length, lower right panel. In each panel the lighter bars represent the non-pregnant women, and the darker bars the pregnant ones.

Fig. 4 displays the distribution of anatomic properties in terms of pregnancy outcome. In each panel the width of the bar is proportional to the total size of the class population. Visual inspection suggests that the greater the differences in segment location and diameters, the greater the likelihood of miscarriage and ectopic pregnancy. The presence or degree of fibrosis has little apparent effect on pregnancy outcome.

**Figure 4.**
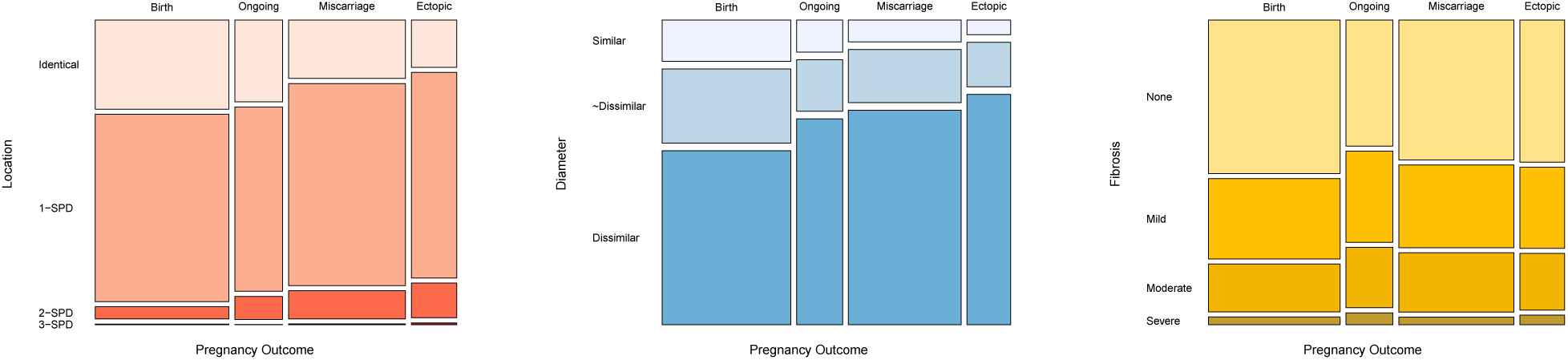
Populations of different pregnancy outcomes in terms of categorical tubal anatomic properties. From left to right: Location, Diameter and Fibrosis properties. In each panel the width of the bar is proportional to the total size of the population.

In what follows, we introduce a more quantitative analysis of all properties simultaneously for their influence on pregnancy and outcome likelihoods.

## 2 Methods

### 2.1 Generalised Additive Models

The simple *linear regression* model, although ubiquitous, falls short when the data to be modelled come from *exponential family* distributions other than the Normal/Gaussian^6–9^. For such problems, there is a solution known as generalised linear models (GLMs). Regression models in the class of GLMs^10^, take a more general form than in ordinary linear regression:

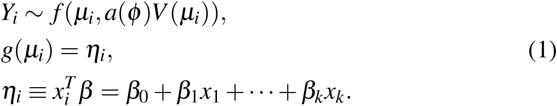

In equation (1), *f* denotes a response variable distribution from the exponential family (EF), *µ*_*i*_ is the response variable mean, *ϕ* is the EF dispersion parameter in the dispersion function *a*(·), *V* (*µ*_*i*_) is the response variable variance function, *η*_*i*_ is the linear predictor, 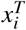 is a vector of explanatory variables (covariates or predictors), *β* is a vector of covariate coefficients, and *g*(·) is the link function, which connects *µ*_*i*_ to *η*_*i*_.

The methodology discussed herein focuses on a particular class of GLMs known as logistic regression, which is suitable for handling Bernoulli (or binomial) distributed data. Bernoulli distribution is a particular case of the more general binomial distribution, Binomial 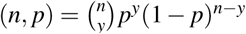, for which *y* is the number of successes (*y* = 1), *n* is the number of trials, and *p* is the probability of success. For the Bernoulli distribution, Bernoulli(*p*) = *p*^*y*^(1 – *p*)^1^*-y*, the number of trials, *n*, is set to 1.

The natural link function for the Bernoulli distribution is known as the logit link, which defines the logistic model:

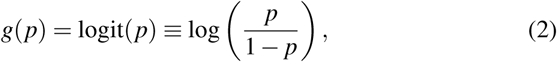

and ensures a bijection between the (-∞, ∞) range of *η*_*i*_, and the (0,1) range of non-trivial probabilities for the Bernoulli *p*.

As for our analysis, we employed an important extension of the GLM methodology known as generalised additive models (GAM)^11^. The GAM model, in our context, extends the linear assumption by assuming the existence of an unknown functional relationship between the expected value of a given property *E*(*y*) and a set of covariates *x*. In other words, *E*(*y*) = *f* (*x*) for unknown *f*. GAMs assume that *f* (*x*) = *f*_1_(*x*_1_) + *f*_2_(*x*_2_) + *…* + *f*_*p*_(*x*_*p*_).

Throughout this work, we evaluate the GAM model using the implementations^12, 13^ within the R language^14^.

### 2.2 Cross-Entropy Loss

The cross-entropy loss, *H*_*p*_, measures the performance of a classification model whose output is a probability value between 0 and 1^15^. The *H*_*p*_ increases as the predicted probability diverges from the actual label. Hence, the lower the *H*_*p*_, the better is the model’s predictive capabilities.

In the case of multi-classification problems, with *κ* classes, we estimate a loss for each class label per observation and sum the result:

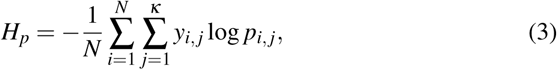

where *κ* is the number of classes (birth, miscarriage, ectopic, outgoing), y is a binary indicator if class label *j* is the correct classification for a given observation *i, p* is the predicted probability from the GAM model for a given class *j*. In the particular case of a binary classification, the equation simplifies as:

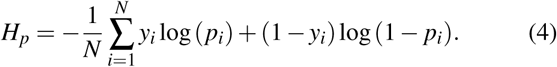

## 3 Results

This section describes the relationships between pregnancy occurrence and outcomes against women’s age and tubal anatomic properties. For that we utilise partial dependence plots (PDP)^16, 17^. PDPs are useful to visualise the relationship between a subset of the features and the response while accounting for the average effect of the other predictors in the model.

### 3.1 Model fit

Figs. 5 and 6 display the conditional fits for pregnancy likelihood in terms of women’s age and tubal length for each of the categorical anatomic properties. The shape and steepness of the curves are indicators of the predictor’s relative influence.

**Figure 5.**
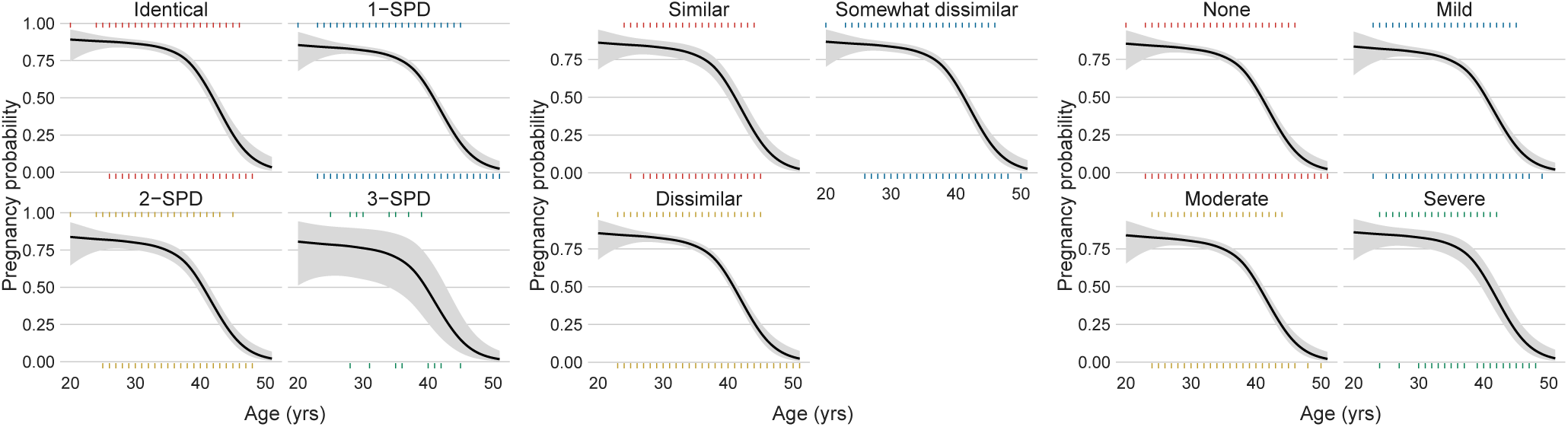
Likelihood of pregnancy by age for each of the observed anatomic properties. From left to right, Location, Diameter, and Fibrosis. The shaded grey areas depict 95% confidence intervals.

**Figure 6.**
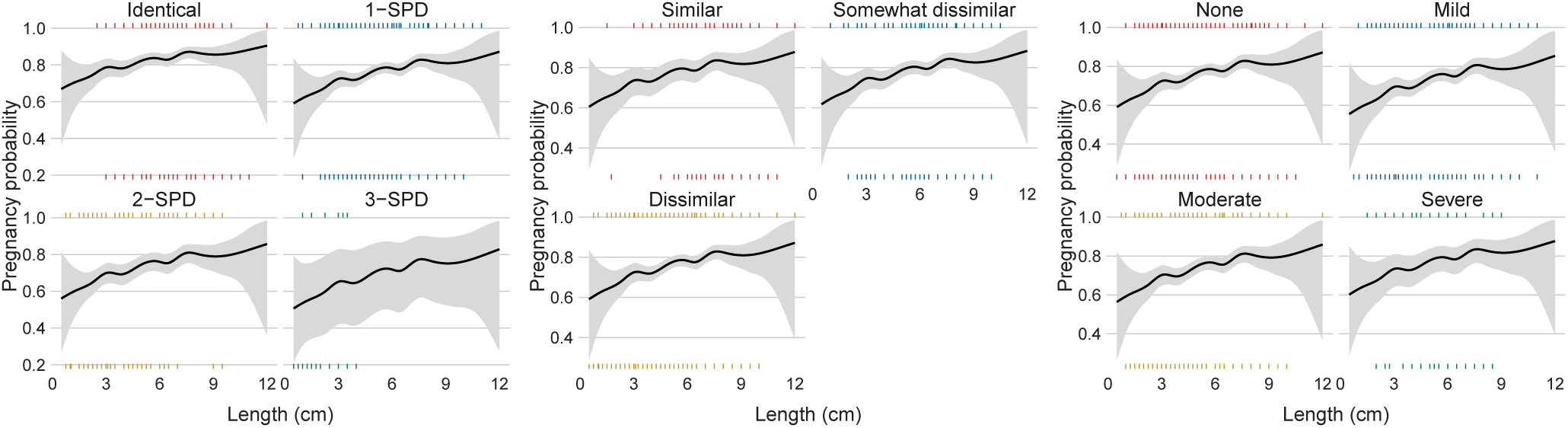
Likelihood of pregnancy by tubal length for each of the observed anatomic properties. The shaded grey areas depict 95% confidence intervals.

Fig. 5 shows the predominant role of age in pregnancy occurrence, which is consistent regardless of women’s different anatomic properties. Pregnancy probability declines steadily as age increases, the rate of decline increasing sharply after ≈ 35 years of age. The higher uncertainty for women with 3-SPD location is due to the small number in this group, as can be seen in Fig. 4.

Tubal length is another good predictor of pregnancy likelihood, as portrayed in Fig. 6. The longer the fallopian tube, the higher the pregnancy odds. This trend is also independent of other properties.

To summarise, age and tubal length are the major factors determining the odds of pregnancy.

Fig. 7 shows the trends between outcomes likelihood as a function of age (left plot) and tubal length (right plot). The left plot quantifies the well known association between age and birth odds. Birth likelihood peaks by age 30 with the chances dropping sharply in the 40s as the likelihood of miscarriage rises with advancing age. The ectopic and ongoing pregnancy groups show no clear relationship with age. The right panel shows the relationship of tubal length to pregnancy outcome. As tubal length increases, the probability of birth rises dramatically; conversely, the odds of both miscarriage and ectopic pregnancy decline.

**Figure 7.**
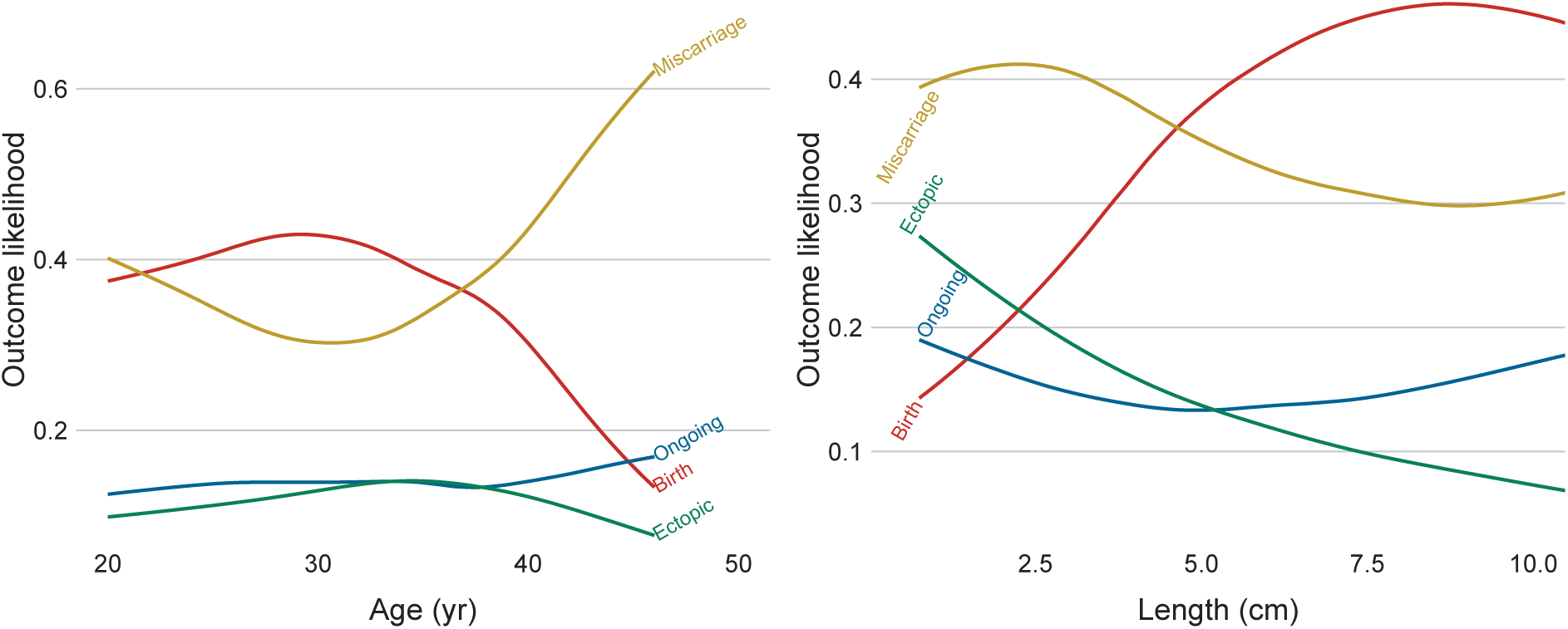
Likelihood of pregnancy outcomes (birth, miscarriage, ectopic, ongoing) by age in the left panel and by tubal length in the right panel from the GAM model.

From Fig. 7, we can conclude that women at ages ≲ 30 yrs and tubal length ≳ 5 cm have the highest odds of a successful pregnancy. On the other hand, women ≳ 40 yrs old and tubal lengths (≲ 5 cm) have the highest risk of miscarriage or ectopic pregnancy.

### 3.2 Feature relevance

A more formal approach to evaluate the importance of the women’s multivariate and interrelated characteristics is to use the cross-entropy loss for assessing the significance of each property in the GAM model after taking the others into account. Specifically, the idea is to quantify the hypothesis that a given woman’s feature has no influence on the probabilistic threshold above which pregnancy or a given pregnancy outcome can occur. The influential rank of each property for pregnancy likelihood is shown in Fig. 8 and for pregnancy outcome likelihood in Fig. 9.

**Figure 8.**
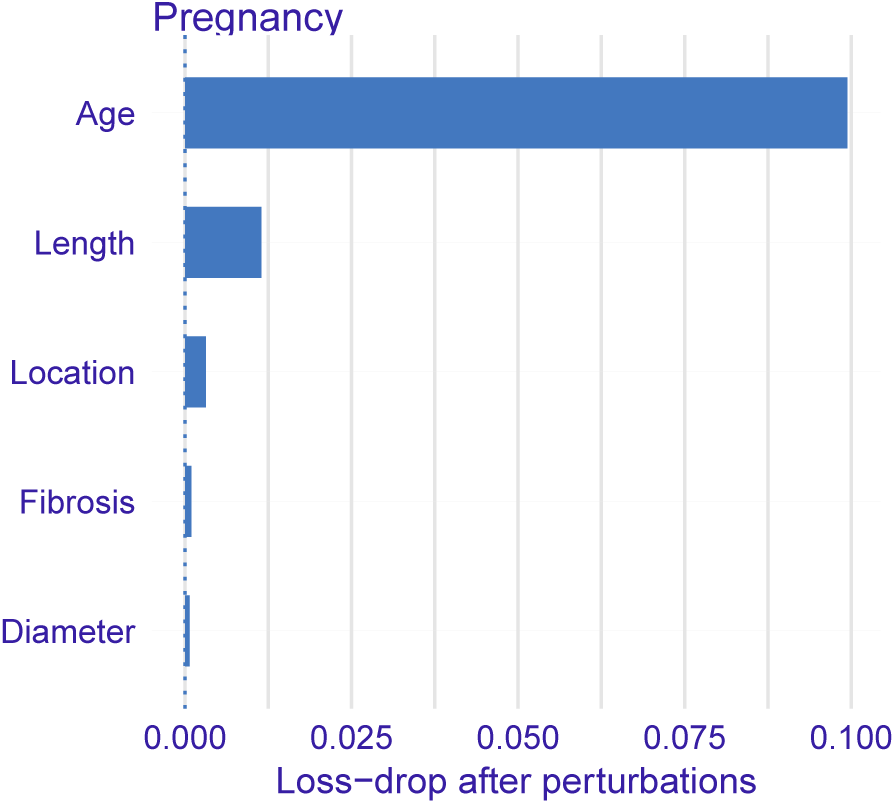
Age and tubal anatomic properties ranked according to their influence on pregnancy likelihood. The x-axis depicts the loss-drop based on the cross-entropy loss function for each feature.

**Figure 9.**
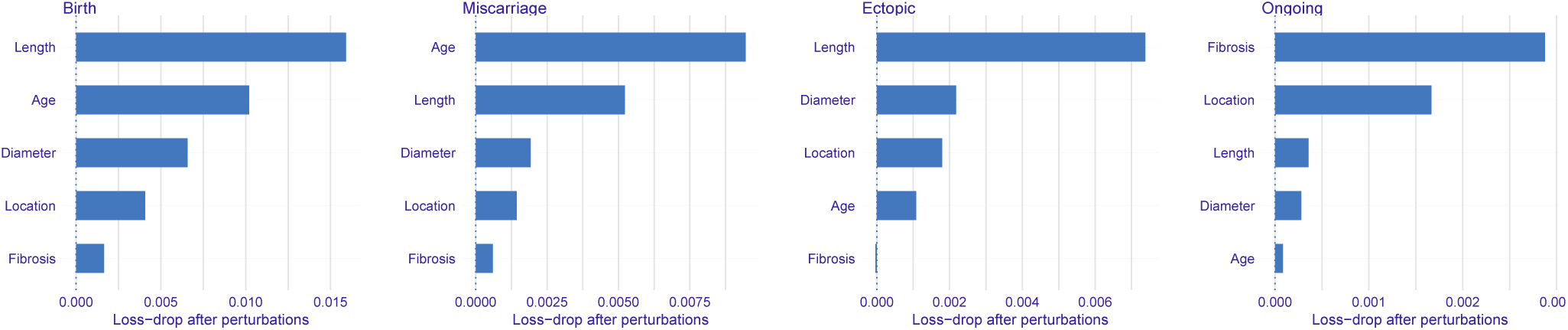
Women’s age and anatomic properties ranked according to their influence on each pregnancy outcome. The x-axis depicts the loss-drop based on the cross-entropy loss function for each feature.

Fig. 8 shows that in predicting pregnancy, age and tubal length are by far the most influential properties, followed by anastomosis location. Diameter differences and fibrosis play less important roles.

Fig. 9 portrays the influential properties for each of the pregnancy outcome categories. Tubal length and age are the most influential factors related to the odds of birth and of miscarriage which are correlated inversely. Tubal length is also the most influential factor for predicting a possible ectopic pregnancy; the shorter the fallopian tube, the higher the odds of this event. The dominant influence of fibrosis in the ongoing pregnancy group is difficult to interpret. In these cases the actual outcome is unknown and there may be an unknown bias in this class.

### 3.3 Model evaluation

This section compares the predictive power and model performance based on anatomic properties against a benchmark model based on sterilisation method. The motivation behind this is to determine if knowing the postoperative anatomic properties improves the model’s capability to predict the likelihood of pregnancy and subsequent outcomes.

The following 3 scenarios were tested: A) age and sterilisation method; B) age and tubal anatomy; C) age, sterilisation method, and tubal anatomy. The performance of each model was evaluated using cross-entropy loss (*H*_*p*_). The results are displayed in table 1.

**Table 1.** Cross-Entropy Loss, *H*_*p*_, for the following scenarios A) age and sterilisation method; B) age and tubal anatomy; C) age, sterilisation method, and tubal anatomy. The middle column refers to pregnancy likelihood (PL) and the right-hand column to outcome likelihood (OL).

| Scenario | $H_p$ (PL) | $H_p$ (OL) |
| --- | --- | --- |
| A | 0.551 | 1.251 |
| B | 0.545 | 1.237 |
| C | 0.544 | 1.232 |

For predicting both pregnancy likelihood and outcome likelihood, Model B (age and tubal anatomy) outperforms Model A (age and sterilisation method). Model C, which encodes the full information, achieves the best performance but only marginally more so than Model B.

## 4 Discussion

Previous studies have identified age at surgery as the primary factor associated with the success of sterilisation reversal. Some studies have examined postoperative tubal length and a few have examined the effect of anastomosis location on pregnancy and birth rates. Patient numbers in most studies have been insufficient to analyse the factors simultaneously, and the results have been conflicting^18^.

The current study population, more than 20 times the mean size of prior studies, permits more in-depth analysis than has been possible in the past. It also examines multiple aspects of postoperative tubal anatomy that have not been reported previously.

Preoperative counselling and obtaining informed consent for tubal anastomosis requires discussing the chances of success and of risks, including possible ectopic pregnancy. To assist with this, the initial report from the Tubal Surgery Database^5^ presented tables of pregnancy and pregnancy outcome rates stratified by age and sterilisation method, both of which are known before surgery.

After tubal reversal surgery, patients usually ask how their particular findings may affect their chances of having a baby. Reframing the question, does tubal anatomic information add predictive power for pregnancy and pregnancy outcome probabilities compared what was known before surgery? That is the question this study was undertaken to answer.

Tubal anatomy observed at reversal surgery necessarily is related to the method of sterilisation. Clips are the least damaging, while coagulation can be extensively damaging to the tubes. Tubal anatomic parameters (length, anastomosis location, segment diameters, fibrosis) are all interrelated. Anatomy often differs between right and left tubes in individual patients. And, it is unknown which of the tubes is involved in a given pregnancy. These issues have not been addressed in previous studies, yet they are important to reflect the realities of clinical medicine. The statistical methodology we have described takes all of these issues into account.

Of the predictive models we tested, the model with anatomic properties gives a better fit to the data than the one without. We believe this model to be robust, based on the consistency of trends in the exploratory analysis, visualisation of the fit, and variable importance analysis.

A limitation of our study is the lack of information about other important factors contributing to fertility, including the male partner; frequency and timing of intercourse; and ovulation history. Nevertheless, it probes deeper and with greater statistical power into post tubal reversal fertility than previous studies.

In conclusion, the answer to the question we posed at the beginning of this study is yes. Information about tubal anatomy, most importantly tubal length, acquired at reversal surgery is a better predictor of pregnancy and pregnancy outcome probabilities than what was known beforehand. The clinical implication is clear. Anatomical information is significant during postoperative counselling when discussing prognosis, the primary concern of patients.

## Data Availability

The datasets analysed during the current study are available from the corresponding author on reasonable request.

## Author Contributions

G.S.B. performed most of the surgeries and conducted the follow-up. RSS participated in analyses and interpretation of results. Both authors contributed equally to the preparation of the manuscript.

## Acknowledgements

We thank Christian Iliadis for his encouraging support of this work. RSS acknowledges the support from NASA under the Astrophysics Theory Program Grant 14-ATP14-0007.

